# Honey bulk DNA metagenomic analysis to identify honey biological composition and monitor honey bee pathogens

**DOI:** 10.1101/2024.07.31.605955

**Authors:** Priit Paluoja, Mihkel Vaher, Hindrek Teder, Kaarel Krjutškov, Andres Salumets, Kairi Raime

## Abstract

Honey’s DNA mixture originates from various organismal groups like plants, arthropods, fungi, bacteria, and viruses. Conventional methods like melissopalynological analysis and targeted honey DNA metabarcoding offer a limited view of honey’s biological composition. We conducted a honey bulk DNA metagenomic analysis to characterize the honey’s taxonomic composition and identify honey bee-related pathogens and parasites based on 266 Estonian and 103 foreign honey samples. 70.4% of the DNA in Estonian honey was derived from green plant families like *Brassicaceae*, *Rosaceae*, *Fabaceae*, and *Pinaceae*. Geographical distribution analysis revealed distinct botanical compositions between Estonian mainland and island samples. The bacterial family *Lactobacillaceae* was prevalent overall, reflecting the honey bee microbiota in honey. We detected 12 honey bee pathogens and parasites, including *Paenibacillus larvae* and *Nosema ceranae*. In conclusion, the study underscores the potential of bulk DNA-based and non-targeted metagenomic approaches for monitoring honey bee health, environmental quality, and honey composition, origin, and authenticity.

## Introduction

Honey bees are considered effective large-scale environmental monitors due to their large-scale foraging activities. Their hive products, especially honey, provide a snapshot of the honey bee and honey production environment, containing nectar and pollen DNA from various plant species and DNA sequences from arthropods, fungi, bacteria, and viruses^1,2^. Previous studies focusing on North European honey biological composition based on DNA analysis have identified predominant floral sources such as *Brassica, Trifolium, Malus, Prunus, Fragaria, Medicago, Populus*, and *Solanum*^3,4^, that are widely spread plant genera also in Estonian nature. *Apis mellifera*, as anticipated, is the most commonly detected arthropod species in honey DNA analyses ^1,3^. Additionally, DNA from several other arthropods from honey bee foraging environments, like plant-sucking and honeydew-producing insects, aphids from the order *Hemiptera* have been detected not only from honeydew honey but also from blossom honey^5^. From viruses, mainly *Apis mellifera* filamentous virus (AmFV) has also been identified within honey DNA, which is known to be a ubiquitous dsDNA virus that affects many apiaries throughout Europe and can have mild pathogenetic effects on honey bees^6^. Also, fungi, mostly yeasts, that are known to tolerate high sugar concentration and recognised for their roles in food and beverage production as fermentative agents, such as species from *Zygosaccharomyces*, and fungal pathogens affecting insects or plants, such as *Metarhizium* spp., *Aspergillus* spp., *Nosema (Vairimorpha) ceranae*, *Bettsia alvei*, or *Alternaria alternata*, have also been observed in honey samples^1,3^. Honey DNA has been found to contain common microorganisms from the honey bee gut microbiota, such as *Lactobacillus kunkeei*, as well as pathogens affecting honey bees or plants, and ubiquitous species like *Escherichia coli*^1^. Honey DNA analysis has been used to detect several potential honey bee pathogens, such as *Paenibacillus larvae* – the causative agent of American Foulbrood, *Melissococcus plutonius* – the aetiological agent of European Foulbrood, and *Spiroplasma* species – the agent of the spiroplasmosis^1,3,7^. Screening for pathogens is essential for several reasons. This detection aids in identifying and managing diseases affecting honey bee colonies that are already at an early stage. Colony losses have been linked to pathogens such as *Varroa destructor* or *Nosema ceranae*^8^. Sensitive bulk DNA-based screening allows the detection of infections before visual symptoms appear. For hive health, early detection of pathogens can facilitate timely intervention, potentially saving colonies from devastating diseases. Additionally, understanding the prevalence and spread of pathogens locally and on larger scales can help monitor and manage diseases and invasive honey bee parasites.

Considering the above, the honey composition reflects the surrounding ecological landscape. It helps detect pathogens, map hive health, describe the honey bee foraging and honey production environment, and describe geographical peculiarities, creating a fingerprint of common regional honey and combating food fraud. Traditional methods, such as melissopalynological analysis or DNA metabarcoding, offer a limited view of honey composition. The melissopalynological analysis is restricted to detecting pollen plants, ignoring nectar and honeydew plants and other organisms, including pathogens, that leave DNA traces in honey^9^. DNA metabarcoding expands this scope by targeting a broader range of organisms, but it remains a targeted approach, limited to detecting only targeted taxa based on a few successfully preamplified genomic regions^10^. To use an unbiased approach, we used shotgun metagenomics sequencing of all DNA extracted from honey sample, which describes the complexity of samples containing thousands of distinct species belonging to different kingdoms or phyla^10^. We conducted a thorough all-DNA-sequencing-based metagenomic analysis on 266 Estonian centrifugally-extracted honey samples to map the botanical composition of Estonian honey with geographical distribution analysis. Additionally, we included 103 foreign honey samples for comprehensive honey bee-related pathogens and parasites analysis, as not all honey bee-related pathogens and parasites of interest are present in Estonia.

## Methods

### Honey samples

A total of 264 honey samples were collected from various regions across Estonia to describe the DNA taxonomic composition of Estonian honey (**S1 Fig**). Additionally, two positive control samples from the hives with diagnosed American Foulbrood infection caused by *Paenibacillus larvae* were included, although their specific locations were not disclosed and are therefore included in honey bee pathogen analysis but not in the analysis of Estonian honey DNA botanical composition and geographical distribution. For honey bee pathogen analysis, in addition to the Estonian honey samples, 103 foreign samples were obtained directly from beekeepers, shops, or honey markets (Error! Reference source not found.). All samples were produced during the summers of 2020 to 2022 and collected for analysis between 2020 and 2023. All honey samples were collected from centrifugal extracted honey and not directly from the honeycomb. Centrifugally extracted honey samples contain DNA traces from several honeycombs and several hives in the apiary and provide a more comprehensive DNA taxonomical composition picture of the honey that is sold on a market as well as the honey bees’ foraging, hives’, and honey production environment in an apiary.

### DNA extraction and sequencing

Each honey sample was preheated at 40°C and homogenized by mixing with a clean spoon. 30 g honey was weighed into a 50 ml centrifuge tube and diluted in 25 ml of preheated MilliQ water. After centrifugation at 4000 rpm, the supernatant was removed, and bulk DNA from the pellet was extracted by NucleoSpin Food Mini kit (MACHEREY□NAGEL). The DNA was fragmented down to 150-200 bp fragments by Covaris M220 focused-ultrasonicator (Covaris) and concentrated by NucleoSpin Gel and PCR Clean-up kit (MACHEREY□NAGEL). The quality and quantity of the DNA fragments were assessed on Agilent 2200 TapeStation (Agilent Technologies). Illumina-compatible DNA libraries were prepared using the Celvia CC AS in-house developed FOCUS protocol. Briefly, fragmented 25 µl honey bulk DNA (1 ng/µl) was end-repaired and A-tailed by a specific enzymatic mixture. Short double-stranded and index-labelled DNA adapters were ligated to both ends of pre-treated DNA fragments. The full adapter sequence and sufficient ready-made Illumina-compatible library were ensured by 12-cycle PCR. 36 samples were pooled equimolarly, and the quality and quantity of the pool were assessed on Agilent 2200 TapeStation (Agilent Technologies). The honey bulk DNA pooled library was sequenced using the Illumina NextSeq 500 instrument (Illumina Inc.) and 85 bp single-read protocol.

### Metagenomic analysis

Sequencing read counts ranged from 1 to 27 million, with a median of 13.7 million reads per sample. Sequencing data analysis was managed with Nextflow (23.09.3-edge) and conducted using the computational resources of the High Performance Computing Center of the University of Tartu^11,12^. Sample sequencing lanes were concatenated, assessed for quality using SeqKit stats (2.4.0), and subsequently analysed for taxonomic composition^13^. No additional pre-processing was applied to the FASTQ files.

To classify the taxonomic composition of the Estonian honey by assigning taxonomic labels to sequence reads, we utilized the taxonomic sequence classifier Kraken 2 (2.1.3) with a custom reference database^14^. The minimum hit groups required for classification were set to 3, and the confidence threshold representing a non-probabilistic scoring scheme for taxonomic assignment was set to 0.5. Across 264 Estonian honey samples, the median percentage of classified sequencing reads was 36.5%, ranging from 9.12% to 71.5%, with a standard deviation of 10.1%.

We constructed a comprehensive reference database by combining data from three sources: the NCBI nucleotide (nt) collection, the One Thousand Plants Project (1KP), and the NCBI Sequence Read Archive (SRA)^15–17^. To ensure the database is highly relevant, we focused on species commonly associated with honey. This process involved two steps: identifying gaps in the coverage of known honey plants and analysing unclassified DNA reads from honey samples. For the latter, we used computational tools to match these unclassified sequences with entries in the SRA database^17^. By selecting the most relevant sequences, we enriched the database to better capture the diversity of organisms found in honey. Sequences from the SRA were further processed to ensure high quality (fastp with the parameters cut_front, cut_right, and correction)^18^. Raw sequencing reads were cleaned and assembled into longer fragments (SPAdes 3.14.1), which were then screened to remove contaminants such as human or bacterial DNA^19^. This ensured that the final database included only meaningful sequences for honey DNA analysis. To improve efficiency, we reduced redundancy in the dataset by removing duplicate or overlapping sequences within each species.

To describe and estimate the abundances of honey bee pathogens and parasites in honey DNA on the species level, we used Bracken (2.8) with the read length set to 80, taxonomic level to species, and threshold for the abundance estimation set to 10^20^. We analysed the presence of the following 20 honey bee-related parasites and pathogens: *Acarapis woodi, Acarus siro, Achroia grisella, Aethina tumida, Ascosphaera apis, Bettsia alvei, Braula coeca, Forficula auricularia, Galleria mellonella, Melissococcus plutonius, Nosema apis, Nosema ceranae, Oplostomus fuligineus, Paenibacillus larvae, Senotainia tricuspis, Spiroplasma apis, Spiroplasma melliferum, Tropilaelaps clareae, Tropilaelaps mercedesae,* and *Varroa destructor*.

Finally, read counts were normalised by total read count to ensure comparability across samples.

For the geographical distribution analysis for plant genera, read counts were further normalised by the number of samples analysed within each county. Statistical analysis of the Bracken and Kraken 2 outputs, as well as data visualisation, was conducted in R (4.4.1)^21^.

## Results

Our study presents a metagenomic analysis of honey bulk DNA to identify its taxonomic composition and monitor honey bee pathogens.

### Estonian honey DNA taxonomic composition

In our analysis of the Estonian honey DNA taxonomic composition, we characterised the proportions of bacteria, fungi, animals (Animalia, Metazoa), green plants (Viridiplantae), and viruses (**Fig 1**). As anticipated, most of the DNA was derived from green plants (70.4% ± 0.12), with bacteria constituting a secondary component (22.7% ± 0.07).

**Fig 1.**
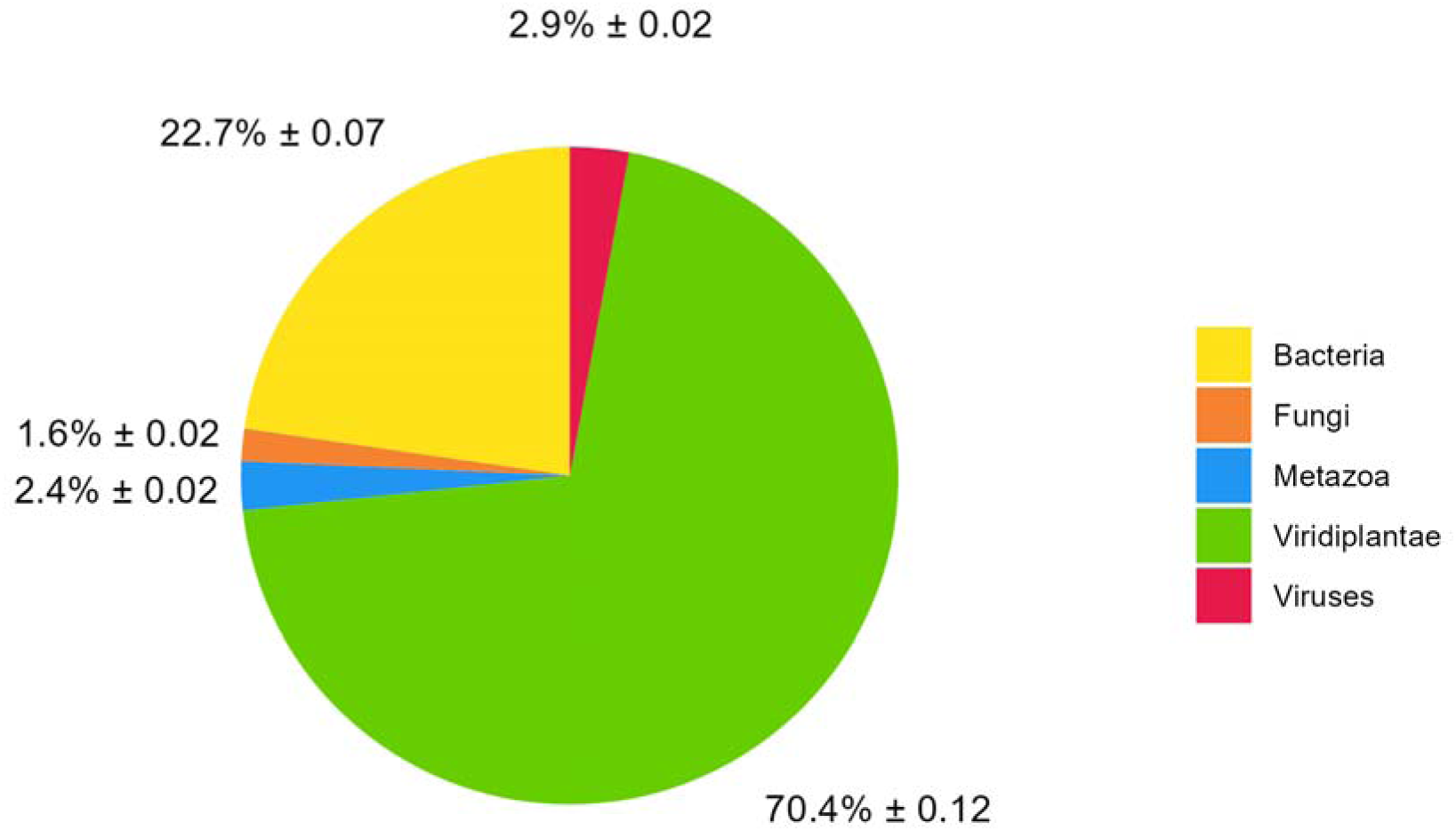
Bulk DNA taxonomic composition of Estonian honey. The symbol “±” represents the standard deviation (s.d.).

Although Viridiplantae dominated the honey composition, the dominant family identified was the bacterial *Lactobacillaceae* (average relative read abundance of 19.5%) (**Fig 2A**). Within the family *Lactobacillaceae*, the prevalent genus was *Apilactobacillus* (**S2 Fig**). Other common bacterial families were *Pseudomonadaceae* (1.7%) and *Erwiniaceae* (1.5%). The top five prevalent families of Viridiplantae in Estonian honey were *Brassicaceae* (19.1%), *Rosaceae* (13.1%), *Fabaceae* (12.0%), *Pinaceae* (9.1%), and *Salicaceae* (7.4%) (**Fig 2A)**. As expected, the genera with the highest abundance were *Brassica*, *Picea*, *Trifolium*, *Rubus*, and *Salix* (**S2 Fig**).

**Fig 2.**
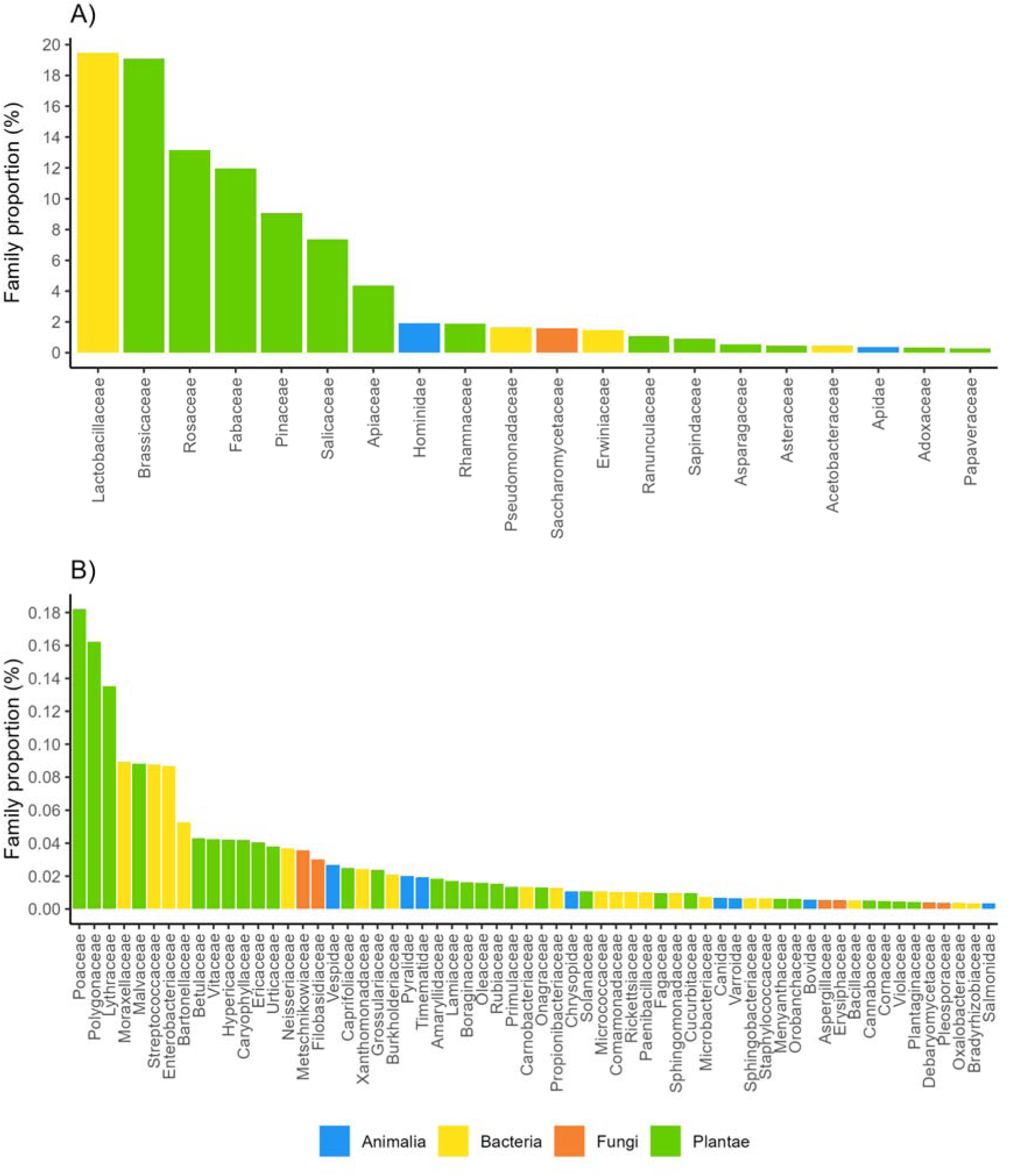
Average relative read abundances of bacteria, fungi, animals (Animalia), and green plants (Plantae) detected in Estonian honey. Panel A shows the relative read abundances for families with an average abundance greater than 0.2% across all Estonian honey samples. Panel B shows families with an average abundance less than or equal to 0.2%.

The prominent Animalia families detected in honey DNA were *Hominidae* and *Apidae*, containing among others human (genus *Homo*), honey bee (genus *Apis*), and bumblebee (genus *Bombus*) DNA (**Fig 2A,B, S2 Fig**). Interestingly, the analysis revealed DNA traces belonging to the mammal families *Canidae* and *Bovidae*, albeit in proportions under 0.2% (**Fig 2A,B, S2 Fig**). Also, DNA from arthropod families containing honey bee or hive parasites or pests can be detected, e.g., *Varroidae*, *Pyralidae, and Vespidae* (**Fig 2B**). The preprominant fungal families detected in honey DNA were *Saccharomycetaceae* and *Metschnikowiaceae*, mainly from yeasts’ genera *Zygosaccharomyces*, *Saccharomyces,* and *Metschnikowia* (**S2 Fig**). Viral DNA was predominantly from the *Apis mellifera filamentous virus (***S2 Fig)**.

### Estonian honey bulk DNA botanical composition and geographical distribution of plant genera

We investigated the geographical distribution of different plant genera of the family Viridiplantae in Estonian honey samples, based on their average relative sequencing read abundances (**Fig 3**). The most widely distributed genus in Estonian honey DNA was *Brassica*, as confirmed by **Fig 2**, where the family *Brassicaceae* was the most common Viridiplantae. While *Brassica* was common and contributed in most areas, there were exceptions. For example, *Brassica* was not dominant on Estonian islands like Vilsandi, Ruhnu, Muhu, Kihnu, and Vormsi (**Fig 3**). Additionally, the islands had different prevalent plant genera compared to the Estonian mainland, such as *Frangula*, *Geum, Rhamnus*, and a considerable proportion of other plant genera (categorised as “Other”) (**Fig 3**). From north to south, the mainland featured common plant genera such as *Brassica, Picea, Trifolium, Salix* and *Rubus*. From east to west, there was an increase in *Rhamnus* and *Frangula* prevalence. Other plant genera, such as *Aegopodium*, *Vicia*, and *Melilotus*, were also prevalent in Estonian honey.

**Fig 3.**
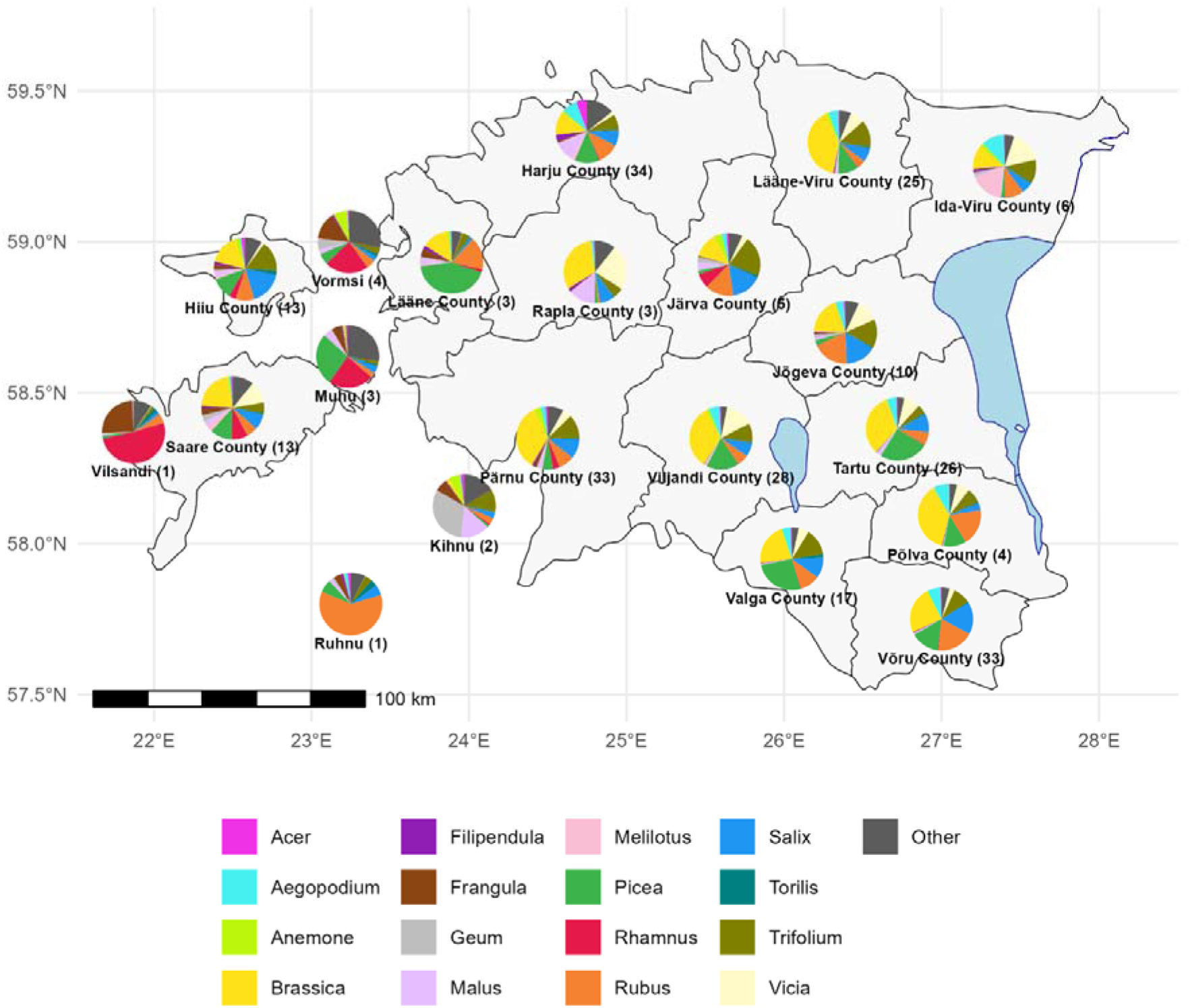
Honey bulk DNA botanical composition and geographical distribution of 16 most prevalent plant genera across Estonia. Each pie chart represents a county, showing the proportional composition of plant genera identified in honey samples. The color-coded legend indicates the corresponding plant genera.

### Honey bee pathogens and parasites in honey bulk DNA

In addition to plants, our methodology also detects DNA traces from animals, fungi and bacteria, including honey bee-related pathogens and parasites. We pre-selected and monitored 20 honey bee pathogens and parasites in Estonian and foreign honey samples (see **Methods**). Specific DNA sequences from 12 pathogens or parasites (out of 20 monitored) were detected in numerous samples with either laboratory-confirmed pathogens, visually confirmed parasites, beekeeper-suspected issues, or samples without confirmation (**Fig 4**, **Fig 5**). For instance, DNA proof from the bacterium *Paenibacillus larvae*, which causes honey bee disease American Foulbrood, was detected in both two laboratory-confirmed control honey samples, each with a fraction of sequencing reads exceeding 2%. In all the samples where the microsporidian parasite *Nosema* sp. was detected, including two samples from the hives suspected of nosematosis, only *Nosema ceranae* was detected but not *Nosema apis*. As expected, DNA traces of *Aethina tumida* (small hive beetle) were only observed in some foreign samples, from USA, Spain and Ghana, as this beetle is not present in Estonia or other European countries. DNA traces from flour mite *Acarus siro* were detected in one Estonian honey sample. The widespread parasitic honey bee mite (*Varroa destructor*) and the greater wax moth (*Galleria mellonella*) were found in many Estonian and foreign apiaries (**Fig 5**). Also, DNA sequences from honey bee pathogens or pests like *Ascosphaera apis* (fungus causing Chalkbrood), *Melissococcus plutonius* (causing European Foulbrood), *Spiroplasma* species (related to spiroplasmosis, May disease), *Bettsia alvei* (causing pollen mold), and even from *Forticula auricularia* (insect, European earwig) were detected in numerous Estonian and foreign honey samples (**Fig 4**, **Fig 5**).

**Fig 4.**
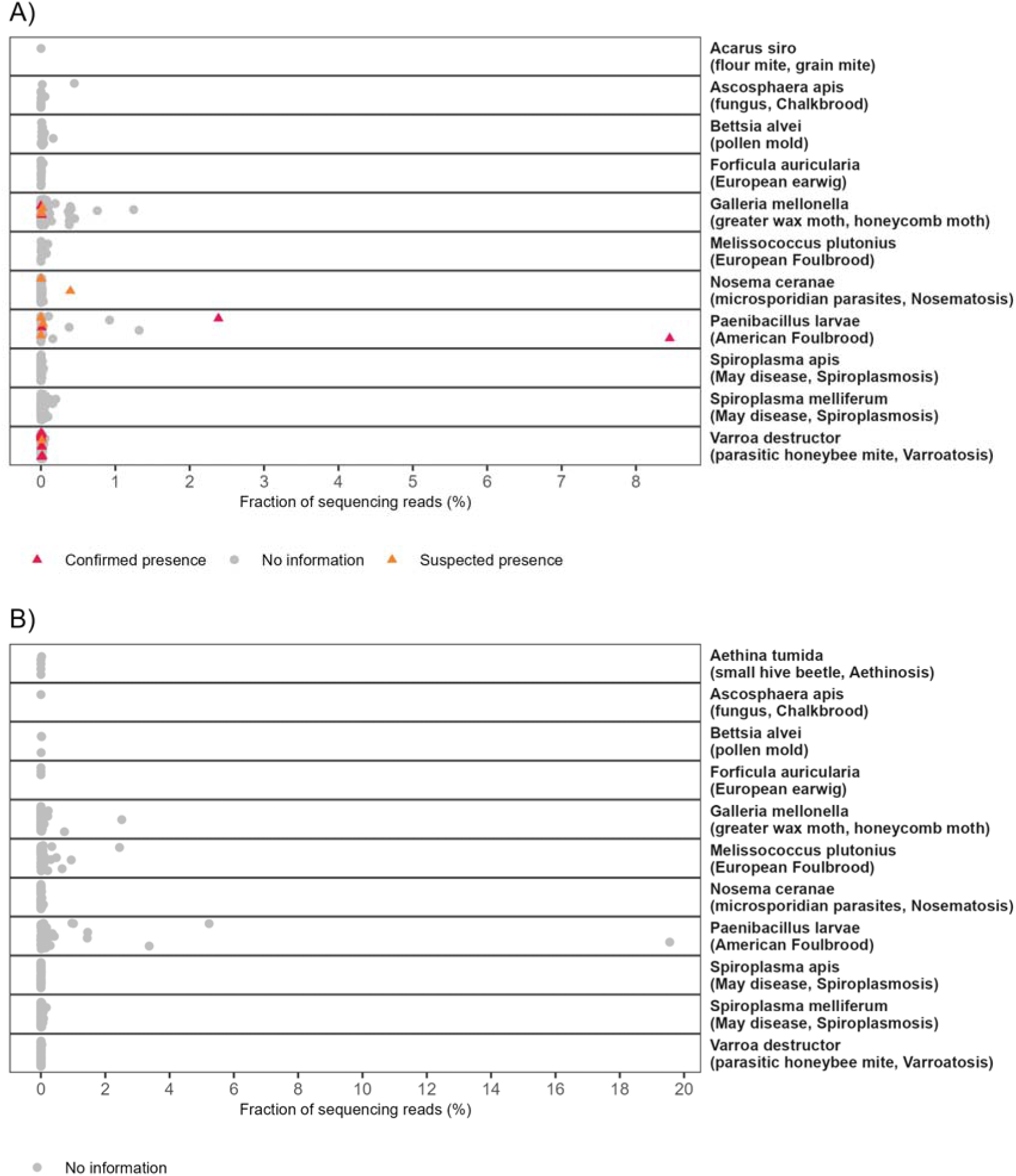
Detection of pathogens and parasites in Estonian (A) and foreign (B) honey samples. Red triangles indicate laboratory-confirmed pathogens or visually confirmed parasites, while orange triangles represent beekeeper-suspected issues. Grey points (“No information”) depict samples with no information. Honey samples that did not yield any sequencing reads assigned to the pathogens listed in the Methods section are excluded from this figure. A fraction close to 0% signifies a very low proportion of sequencing reads assigned to a particular pathogen but indicates presence. Notably, certain pathogens were detected exclusively in either Estonian or foreign honey samples. For example, *Aethina tumida* presence was found only in foreign samples (panel B), whereas *Acarus siro* was detected in only one Estonian sample (panel A). Sequencing reads originating from *Acarapis woodi* were not detected in any of the samples analysed.

**Fig 5.**
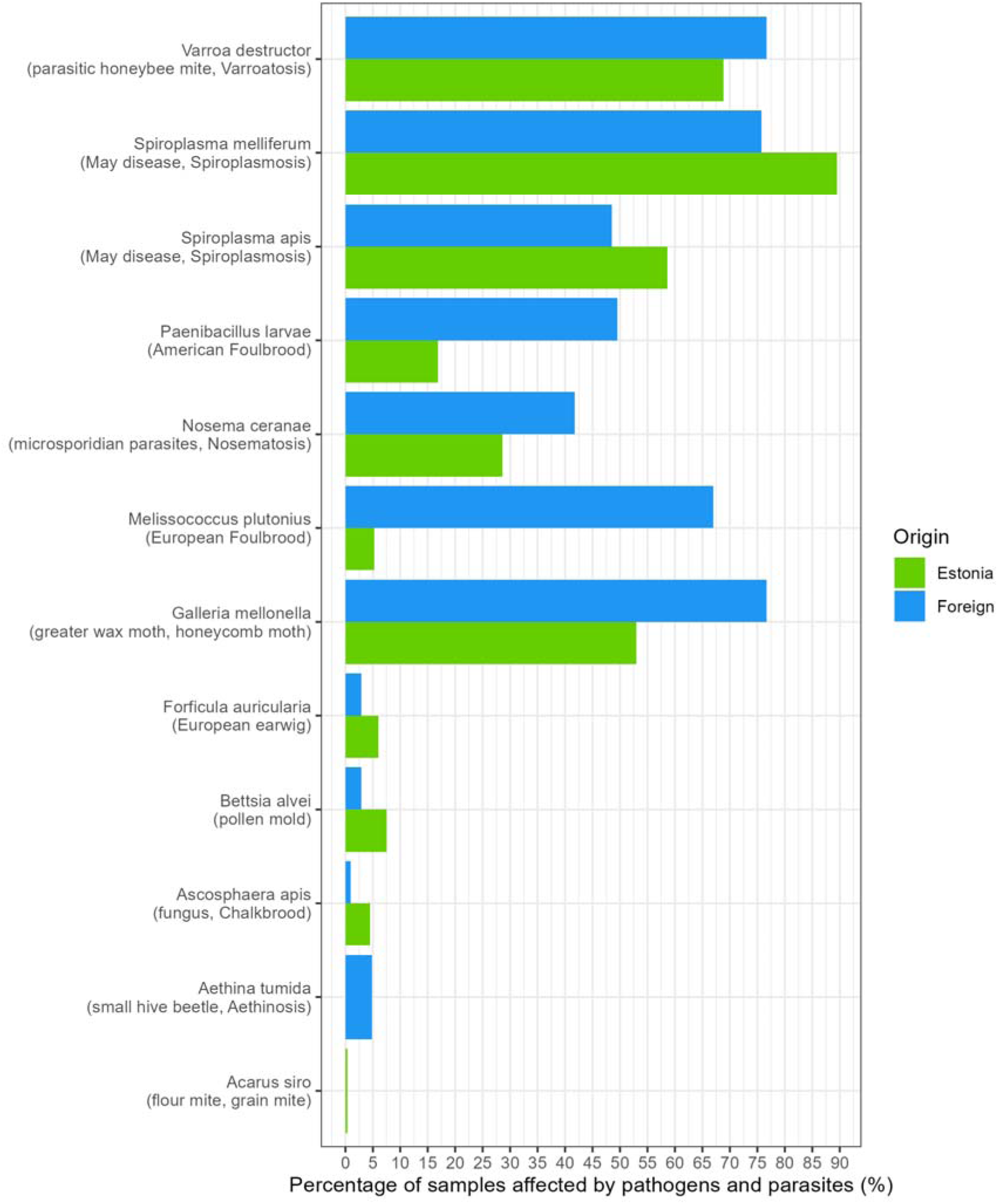
Comparison of the percentage of honey samples from Estonia and foreign origins affected by screened pathogens and parasites.

## Discussion

The honey bulk DNA metagenomic analysis provides a more unbiased and less restricted overview of honey’s biological composition compared to the targeted DNA-based approaches. Unlike the DNA metabarcoding method, which targets limited selected gene(s) of the specific organism(s), the honey bulk DNA approach provides a comprehensive overview of honey botanical, microbial, fungal, viral, and animal (including entomological) diversity, including honey bee pathogens and parasites^22^. We conducted thorough analyses on 266 Estonian and 103 foreign honey samples. Unlike honeycomb-scrapped samples, these samples were collected from centrifugally extracted honey, which contains honey DNA from various honeycombs and hives of the apiaries from different locations. The amount of at least one million metagenomic DNA sequencing reads per honey sample enables us to describe the biological environment of honey bee foraging and honey production. In addition to the DNA only from plant pollen, this method analyses all DNA traces in the sample, including cell-free DNA, which allows us to detect pollen, nectar and honeydew plants as well as DNA from other organisms.

We demonstrate that green plants (Viridiplantae) constitute the majority of the DNA content in honey, accounting for 70.4% of the total honey DNA composition, with *Brassicaceae*, *Rosaceae*, *Fabaceae*, *Pinaceae*, and *Salicaceae* being the most common plant families identified in Estonian honey (**Fig 1**, **Fig 2**). The most common plant genera were expectedly *Brassica*, *Picea*, *Trifolium*, *Rubus*, and *Salix* (**S2 Fig**). These results concord with the observations made for the composition of honey pollen plants in Estonia^23^, indicating that a significant part of plant DNA in honey may originate from plant pollen.

Interestingly, the most predominant genus detected in honey DNA based on the amount of sequencing reads was not from the plants but the bacterial genus *Apilactobacillus*, aligning with its known association with honey bee microbiota (**S2 Fig**), as also shown by the past study^3^. Although in much lower proportions, also other notable bacterial families, like *Pseudomonadaceae* and *Erwiniaceae* (1.7% and 1.5%, respectively), were detected, both of which include species known for their roles in various ecological functions and interactions with plants and insects (**Fig 2)**^24^. These findings demonstrate that the taxonomic diversity of plant genera in honey DNA surpasses that of bacterial genera. Moreover, bacterial taxa are often represented by a few dominant families, while plant DNA is more evenly distributed across numerous genera.

As expected, the most common Animalia families detected in honey DNA were the mammal’s family *Hominidae* and the arthropods’ family *Apidae*, containing mostly human (genus *Homo*), honey bee (genus *Apis*), and bumblebee (genus *Bombus*) DNA from honey bee foraging and honey production environment. Interestingly, DNA from arthropod families containing common honey bee or hive parasites or pests from the honey bee or honey production environment can be detected in honey DNA, e.g., *Varroidae*, *Pyralidae, and Vespidae* (**Fig 2B**). The family *Vespidae* includes species detrimental to honey bees, such as hornets. Although hornet DNA detected in our samples was mainly from the European hornet *Vespa crabro*, this finding could be valuable when searching methods for monitoring and early detection of the Asian hornet (*Vespa velutina*), a species known to be devastating for honey bee populations in warmer areas of Europe, but not yet detected in Estonia ^25^. The widespread parasitic honey bee mite (*Varroa destructor*) from the arthropod family *Varroidae* and the greater wax moth (*Galleria mellonella*) from the family *Pyralidae* were detected both in many Estonian and foreign honey samples (**Fig 4**, **Fig 5)**^26,27^.

In contrast, the small hive beetle (*Aethina tumida*), known to cause colony collapses in weak colonies, was only found in three samples, according to the label originating from the US, Spain, Ghana, and two honey blends of undetermined geographical origins (**Fig 4)**^28^. Importantly, *Aethina tumida*, known to be absent in Estonia, was not detected in any Estonian honey samples (**Fig 4)**. This approach demonstrates that the honey bulk DNA metagenomic analysis could be a valuable screening tool to monitor agriculturally significant honey bee parasites’ prevalence and geographical distribution.

Our analysis revealed the presence of several other honey bee-related pathogens and parasites (**Fig 4)**. Notably, the bacteria species *Paenibacillus larvae*, which is known to cause American Foulbrood disease in honey bees, was detected in several samples, including two positive control honey samples from the hives that were confirmed to have American Foulbrood disease^29^. In both control samples, a substantial proportion of sequencing reads were attributed to *Paenibacillus larvae* (**Fig 4**, 8.5% and 2.4%). Also, DNA traces from honey bee pathogens or parasites like *Ascosphaera apis* (fungus causing Chalkbrood), *Melissococcus plutonius* (causing European Foulbrood), *Nosema ceranae* (microsporidian parasite, causing Nosematosis), *Spiroplasma* species (related to spiroplasmosis, May disease), *Bettsia alvei* (causing pollen mold), and even from *Forticula auricularia* (insect, European earwig) were detected in several Estonian and foreign honey samples. We did not detect DNA of the following honey bee pathogens or parasites in any analysed Estonian or foreign honey sample: *Acarapis woodi* (parasitic honey bee mite, causes acarapiosis), *Achroia grisella* (lesser wax moth), *Braula coeca* (Braula fly, bee louse), *Oplostomus fuligineus* (large African hive beetle), *Senotainia tricuspis* (fly, causes senotainiosis), *Tropilaelaps clareae* (parasitic honey bee mite, causes tropilaelapsosis) and *Tropilaelaps mercedesae* (parasitic honey bee mite, causes tropilaelapsosis). This might be because these important honey bee pathogens and parasites species are not widespread worldwide, and none of these have been seen in Estonian apiaries yet. We also did not detect the microsporidian parasite *Nosema apis* in our samples, even though it has been identified as the primary *Nosema* species responsible for Nosematosis in Estonia^30^. Research has shown that *N. ceranae* has replaced *N. apis* in many countries including Italy, Argentina or even northern countries such as Lithuania^30–34^. Essentially, *N. ceranae* has spread rapidly worldwide^30^. Therefore, it is possible that *N. ceranae* has also replaced *N. apis* in Estonia by now.

Interestingly, we even detected trace amounts of DNA sequences from mammals, probably originating from domestic or pest animals, possibly due to the contamination from the honey bee foraging, honeycombś storage, hove or honey production environment. For example, honey bees often collect brackish water enriched with mineral salts, which could be contaminated by mammal excreta and DNA (*Canidae* and *Bovidae,* **Fig 2**) ^35^. This result shows the sensitivity of DNA analysis and indicates the possible DNA transfer through honey bees’ diet. This is in accordance with the study that has demonstrated the presence of DNA from plant-sucking insects in honey DNA that produce the sticky excretion collected by honey bees^5^. DNA contamination from pest animals, such as mice representing <0.2% of sequencing reads, may result from their contact with the honeycombs or the hive environment.

The fungal community was primarily represented by *Saccharomycetaceae* and *Metschnikowiaceae*, families of yeasts, mainly genera *Zygosaccharomyces*, *Saccharomyces*, and *Metschnikowia*, commonly involved in fermentation processes (**S2 Fig**). The presence of *Saccharomycetaceae* has also been detected in previous honey-related studies^1,3,36^. We also detected viral DNA, predominantly from the *Apis mellifera filamentous virus* (**S2 Fig**), which is known to infect honey bees but is little to no pathogenic and has been detected in past studies^6,37^. The difference between our finding of 2.9% sequencing reads assigned to DNA viruses and the 40.2% (± 30.0%), as reported by Wirta *et al.* ^3^, can be explained by differences in the reference database and the number of samples analysed (**Fig 1**). However, most honey bee disease-causing, such as Deformed Wing Virus and many other, are RNA viruses that need RNA-based screening methods^38^.

We also investigated Estonian honey DNA botanical composition and geographical distribution of most prevalent plant genera (**Fig 3**). Consistent with previous findings, we also observed frequent occurrences of *Brassica*, *Malus,* and *Trifolium*, aligning with previous records from North European honey (**Fig 3)**^3,23,39^. Interestingly, we observed distinct differences in the plant genera compositions between the Estonian mainland and the islands, with the islands showing a higher proportion of *Frangula* and species categorised as “Other” compared to the mainland (**Fig 3)**. In the honey DNA samples from small islands in Estonia, the proportion of *Brassica* was substantially lower compared to the other regions. This could be explained by the lack of large agricultural fields on small islands in Estonia. Furthermore, the diverse DNA taxonomical composition of honey creates a unique fingerprint for every honey sample containing hundreds of different species of plants, bacteria, fungi, animals/arthropods/ and other organisms. Therefore, we hypothesise that metagenomic analysis of all extracted DNA could be utilised to analyse the authenticity and geographical origin of honey (**Fig 2**, **Fig 3)**.

Metagenomic analysis of honey DNA presents inherent challenges, primarily because the accuracy of the results heavily relies on the public reference database used for analysis, as also pointed out by other researchers ^40^. If a genus is absent from the database, it can introduce biases and potentially reduce the accuracy of the analysis ^40^. As comprehensive databases for plants are still under development and there is a predominance of complete genome sequences for bacteria and viruses in existing databases, we created a custom Kraken 2 reference database in our study (including partial genome sequences), with the extended numbers of honey-related plants. Our Kraken 2 reference database was sourced from three main collections: NCBI nt collection, The One Thousand Plants Project, and NCBI’s Sequence Read Archive ^15–17^. This approach enables the detection of an increased number of plants in honey DNA. In addition, the majority of foreign honey samples used in the pathogen analysis in this study were acquired from shops, the contents of the honey jars were not validated, and we had to rely on the label information. However, as we were using foreign honey samples only for pathogen analysis in this study, the accuracy of the label did not affect the proof-of-concept of detecting known pathogens in honey samples.

In conclusion, our metagenomic analysis of honey DNA provided a detailed and comprehensive overview of its biological composition, highlighting its significant diversity of botanical composition, and honey bee-related pathogens and parasites. This study mapped the botanical composition of Estonian honey with geographical distribution of different plant genera in Estonia, and conducted honey bee-related pathogen analysis, underscoring the potential of all DNA sequencing-based metagenomic approaches not only for describing the botanical composition of honey, monitoring honey bee health and apiary environment but also for identifying authenticity and origin of honey by using untargeted analysis of all DNA sequences extracted from honey.

## Data availability

The data generated during this study is available in the Sequence Read Archive (SRA) repository under BioProject PRJNA1135913 (https://www.ncbi.nlm.nih.gov/sra/PRJNA1135913).

## Supporting information

**S1 Fig.**
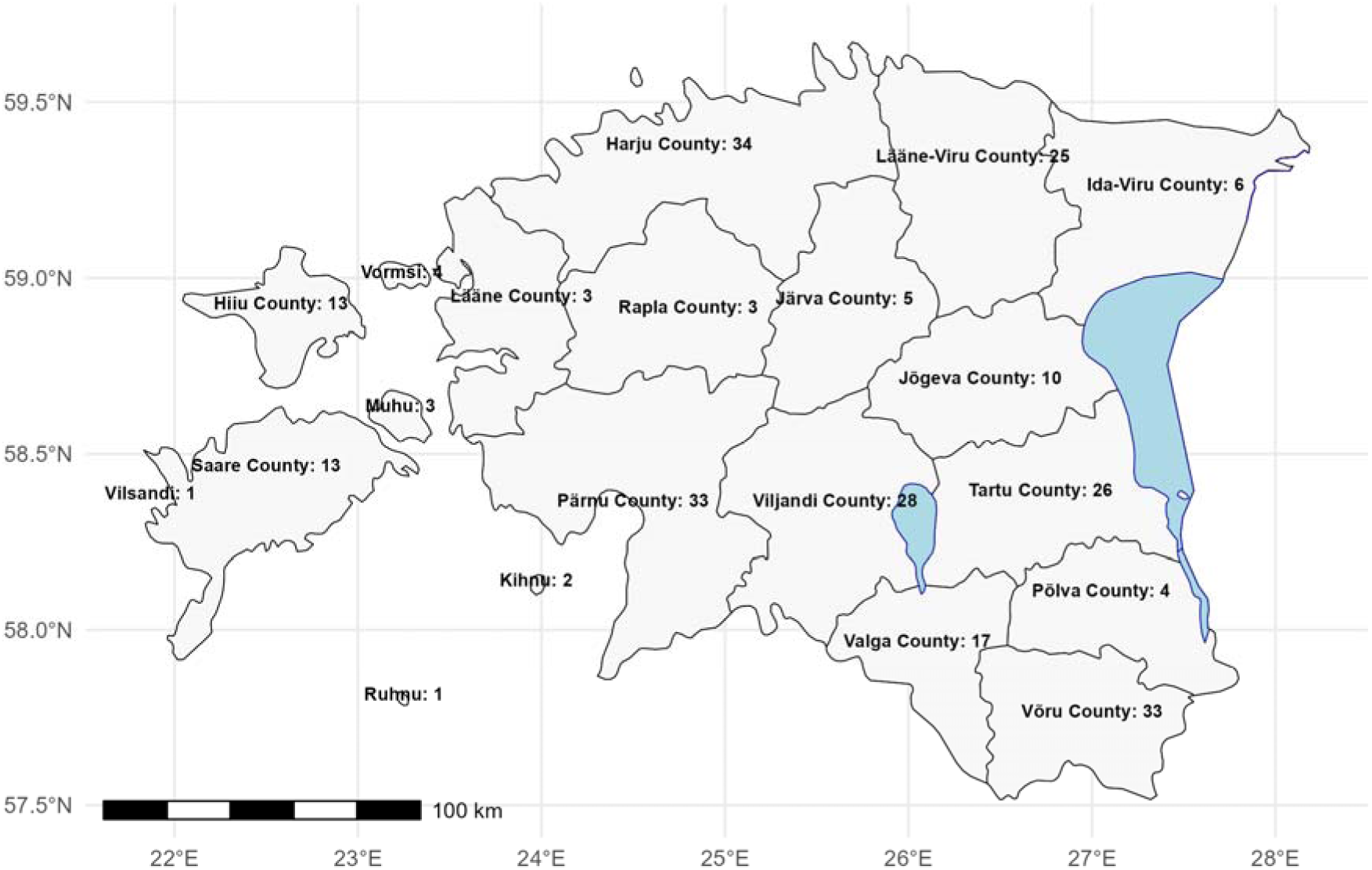
Geographical distribution of 264 Estonian honey samples used in the study.

**S2 Fig.**
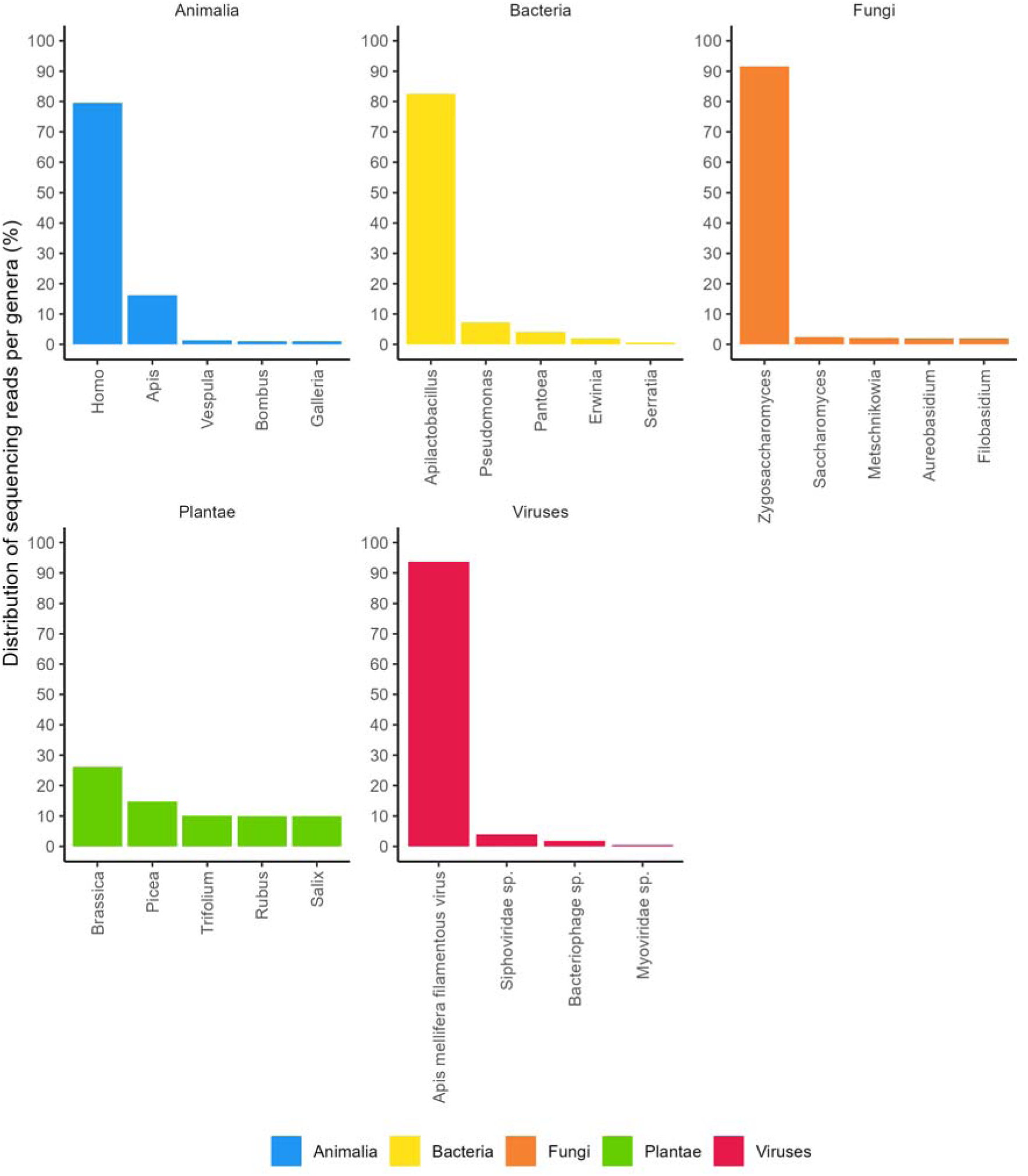
Common genera of Bacteria, Fungi, Animalia, Plantae, and Viruses from the DNA of Estonian honey.

**Table S1.** Origins of 103 foreign honey samples utilised in the pathogen analysis.

| Origin | N |
| --- | --- |
| Austria | 1 |
| Bulgaria | 9 |
| China | 10 |
| Denmark | 2 |
| England | 2 |
| Finland | 3 |
| Germany | 1 |
| Ghana | 1 |
| Greece (including Rhodes Island) | 4 |
| India | 4 |
| Italy | 1 |
| Kazakhstan | 1 |
| Latvia | 4 |
| Mix of EU and non-EU honey | 18 |
| Montenegro | 1 |
| Non-EU honey (including Ukraine, Central and South American honey) | 7 |
| Poland | 3 |
| Portugal | 1 |
| Saudi Arabia | 1 |
| Scotland | 3 |
| Spain | 2 |
| Switzerland | 2 |
| Ukraine | 17 |
| United Arab Emirates | 1 |
| Unspecified EU honey | 2 |
| USA | 1 |
| Yemen | 1 |

